# Establishing Prokaryotic Expression System of Angiotensin-Converting Enzyme 2 (ACE2) gene in pigs

**DOI:** 10.1101/2020.03.12.988634

**Authors:** Hang Xiao, Xin-Tian Nie, Xiao-Xia Ji, Shu-ping Yan, Bin Zhu, Yuan-Shu Zhang

**Affiliations:** Key Laboratory of Animal Physiology and Biochemistry, Ministry of Agriculture, Nanjing Agricultural University 210095; Xintian Nie, Key Laboratory of Intelligent Agricultural equipment in Jiangsu Province, Nanjing Agricultural University (China). E-Mail

**Keywords:** Angiotensin Converting Enzyme 2, Pig, Gene cloning, Bioinformatics analysis, Expression

## Abstract

In this paper, ACE2 gene of pigs was cloned and the purified protein was obtained via the prokaryotic expression system. Polyclonal antibody of high titer and sensitivity was obtained using Wastar rats immunization method and is then used to determine of the expression of ACE2 using immunohistochemistry. The sequence of ACE2 in pigs covered 2418 nucleotides and coded 805 amino acid (aa) residues. Sequence homology analysis showed that the ACE2 sequence in pigs is highly conserved among species at the nucleotide and amino acid levels. Genetic evolution analysis revealed that ACE2 gene in pigs has the shortest genetic distance with that in goats while residing in a totally different branch from that in zebra fishes. Analysis of protein structure predicted that ACE2 protein is a transmembrane secreted protein with high hydrophilicity, containing a signal peptide sequences locating between 1aa to 17aa. The ACE2 fusion protein expressed (under the induction with 1.0 mmol/L IPTG for 10 h) was of approximately 100 kDa and mainly existed in inclusion body. Wastar rats immunization showed that the titer of the anti-ACE2 antiserum in rats was 1: 3200. Western blot showed that the antibody binds specifically. Immunohistochemistry showed that the ACE2 protein was expressed in all major tissues of pigs. It is the first time that polyclonal antibody of ACE2 in pigs was obtained and the expression of ACE2 was confirmed. These results will provide a basis for investigating on ACE2’s biological activity in pigs.

## 1 INTRODUCTION

The renin–angiotensin system (RAS) is an importance regulator of cardiovascular and renal function, both under physiological and pathological conditions(Tipnis et al., 2000). Angiotensin converting enzyme 2 (ACE2) was discovered as a new member of the RAS system in 2000. Earlier studies showed that ACE2 in rats or humans was highly expressed in the kidney, heart and testicle tissues, and was later confirmed existence in the lungs, brain, and the intestines and other organs(Lambert et al., 2010; Pei et al., 2010; Briones et al., 2010; Imai et al., 2005) The ACE2 genes in human, rat and mouse have been located on the X chromosome. The ACE2 protein main structure consists of an extracellular N terminal catalytic domain, a transmembrane region and a C terminal anchored on the cell membrane(Danilczyk et al., 2006). The metalloproteinases, such as ADAM metal protease domain 17 (ADAM17), can hydrolyze the extracellular domain (which is of enzyme activity) to produce active soluble ACE2 form, which has carboxypeptidase activity and participates in the hydrolysis of AngI, AngII and other peptides. Among all, the main function of it is to hydrolyze the phenylalanine at the C-end of the AngII to produce Ang (1-7). Ang (1-7) is later combined with the Mas receptor and plays various protective functions, which antihypertensive activities and exhibits some biological actions that are quite distinct from Ang II(Hashimoto et al,. 2012; Wang et al,. 2016).

So far, the research on ACE2 has been focused on rodents and human beings. The research gaps in other animals, especially production animals, such as cattle, sheep, pigs still need to be filled. Particularly, GenBank still lacks the information of ACE2 gene in pigs. Recently, Zhang has reported a segment of the ACE2 gene sequence in Chinese Jiangsu local Hybrid Pigs(Zhang et al,. 2012). However the size of the gene fragment is only 641 bp.

Therefore, this study was aimed at determining the whole cDNA sequence of porcine ACE2 gene using gene cloning and other molecular biology techniques and establishing a prokaryotic expression system of ACE2 in pigs. The porcine ACE2 polyclonal antibody was obtained and the expression distribution of ACE2 was determined. We hope to be able to provide the whole sequence of ACE2 gene and other related information for Genbank.

## 2 MATERIALS AND METHODS

### 2.1 Ethics Statement

All animal procedures were approved by the Institutional Animal Care and Use Committee of Nanjing Agricultural University. The protocols were reviewed and approved, and the project number 2011CB100802 was assigned. The slaughter and sampling procedures strictly followed the ‘Guidelines on Ethical Treatment of Experimental Animals’ (2006) no. 398 established by the Ministry of Science and Technology, China and the ‘Regulation regarding the Management and Treatment of Experimental Animals’ (2008) no. 45 set by the Jiangsu Provincial People’s Government.

### 2.2 Samples collection

The Healthy newborn piglets were terminally anesthetized via an intraperitoneal (ip.) injection of 20% urethane (5 mL/kg body weight), were decapitated within 15 min after anesthesia took effect (i.e. animals lost conciseness and whole body muscle relaxation was observed), and all tissues including heart, liver, hung, kindey, stomach and soon were collected and frozen immediately in liquid nitrogen and then stored at −80°C, Part of tissues were fixed immediately with 4% buffered paraformaldehyde for 24 hours, then transferred to 20% sucrose solution for 3 h.

### 2.3 Primer design

Primers for the ACE2 and β-actin genes were designed with Primer 5.0 software and were synthesized by Dingguo, Beijing, China based on their respective porcine sequences to produce an amplification product. which the primers for β-actin (an I nternal control) were forward 5’-GATCTGGCACCACACCTTCT-3’, and reverse 5’-CCAGAGGCATACAGGGACAG3’; and the primers for ACE2 were forward 5’ GGATCCATGTCAGGCTCTTTCTGGCT-3’, and reverse 5’-GAGCTCCTAAAACGAAGTCTGAATGTCATCGC-3’; The expected size of the porcine ACE2 gene product was 2418 bp (GenBank accession no. NM_001123070).

### 2.4 RNA extraction and RT-PCR

Total RNA was extracted by using the RNeasy mini kit (Qiagen, Valencia CA), and RNA concentrations were determined by a ultraviolet colorimetry (ThermoScientific, Wilmington, DE). One microgram of total RNA was reverse transcribed to cDNA using two step reverse transcription inversion following the manufacturer’s protocol. ACE2 mRNA expression were amplified by PCR with β-actin as an internal control (Bt03279175); The amplification conditions of PCR reaction were 95°C for 5 min, 32 cycles of denaturation at 98 °C for 10 sec, annealing at 57°C for 15 sec, extension at 72°C for 30 sec, and finally an additional extension step at 72°C for 10 min. The amplified products were identified by 1% agarose gel electrophoresis.

### 2.5 Purification, cloning and sequence analysis of PCR products

#### 2.5.1 Purification of PCR products

The target gene from PCR was recovered by a gel Recovery Kit (Axygen, USA), and the PCR products recovered were connected with pMD-19T carrier(Takara, Janpan), then converted to DH5 alpha receptive cells (Vazyme, China), and evenly coated into the LB agar plate containing 0.1 mM ampilin for single monoclonal colony culture. After 18 h, positive colony was collected. and the bacteria solution was verified by PCR. The PCR reaction and conditions were the same as described previously.

#### 2.5.2 Cloning of PCR products

Omega Plasmid Extraction Kit was used to extract recombinant plasmid vector. The extracted plasmid was identified by single enzyme digestion (HindIII) and double enzyme digestion (BamHI and SacI) according to the Takara enzyme product instructions, and the enzyme digestion products were identified by 1% agarose gel electrophoresis.

The amino acid sequence was deduced from the Editseq program in DNASTAR software and MegAlign software plotted the genetic evolution tree for affinity analysis. SignalP 4.1 (http://www.cbs.dtu.dk/services/SignalP/) predicted the location and sequence of the protein amino acid as well as the sequence signal peptide of the ACE2 gene encoding. TMHMM Server v.2.0 (http://www.cbs.dtu.dk/services/TMHMM/) predict the transmembrane and transmembrane directions of ACE2 encoded protein and Kyte-Doolittle method predicts the hydrophilic region by DNASTAR software.

### 2.6 Construction and identification of recombinant expression vector

The plasmid of pMD-19T-ACE2 and pET-32a was digested by BamHI, SacI enzyme, the target fragment was recovered and the recombinant protein expression vector pET-32a-ACE2 was constructed used T4 DNA ligase, and transformed into Escherichia coli DH5 alpha. The positive recombinant plasmid were screened for the monoclonal bacterial liquid PCR and the double enzyme digestion.

### 2.7 Induction and expression of fusion protein

The Escherichia coli BL21 (DE3) containing the recombinant plasmid containing pET-32a-ACE2 was inoculated into the liquid medium LB containing Amp (100 µg/mL), 37°C, 180 rpm/min until OD_600_ 0.6∼0.8. then The 1.0 mmol/L of IPTG was added to the Escherichia coli BL21 (DE3), and collecting 1 mL bacteria solution after inducting 2, 4, 6, 8, 10 h and overnight respectively. After centrifuging in 8000 rpm/min for 15 min, these bacteria solution were detected by SDS-PAGE.

### 2.8 Identification and purification of the expression of fusion protein

The IPTG stimulation time of high expression was selected according to the results of the former. The bacterial solution was centrifuged in 8000 rpm/min for 15 min, and the bacterial sedimentation was collected and suspended used PBS. The bacterial supernatant and precipitate after ultrasonic breakage (ice bath conditions) were purified, and precipitated by SDS-PAGE. Sure. The fusion protein was purified by staining with reference to Gao Shen Yang(Gao et al,.2010), and the size and purity of the purified protein were detected by SDS-PAGE.

### 2.9 Preparation of anti ACE2 polyclonal antibody

Male New Zealand White rabbits were immunized by subcutaneous injec-tion to gain ACE2 antibody. Briefly, the ACE2 antigen mixed 1:1 with Freund’s complete adjuvant was injected into rab-bits, and booster injections were administered once at10-days intervals with the same amount of antigen mixed 1:1 with incomplete Freund’s adjuvant. Serum was collected from heart method and stored at −80°C.

### 2.10 Extraction of total protein in pig and different species tissues

The protein of heart, spleen, kidney, liver, duodenum, liver of pig, and also liver and kidney tissue of chicken were extracted from Roe. The protein concentration of each tissue was measured by BCA kit(Beyotime Biotechnology, China).

### 2.11 Indirect ELISA method for determination of polyclonal antibody titer of pig ACE2

The serum and pre immune serum of the rat were diluted by 1:50, 1:100, 1:200, 1:400, 1:800, 1:1600, 1:3200, 1:6400, and then incubated in 96 well plate. After washing, Sheep anti rat marked by HRP were added. Finally, TMB was used as the substrate. The OD_450_ value of each well was read on the enzyme standard analyzer and pre immunized serum. As a negative control, the OD_450_ value was N, the serum OD_450_ was P after immunization, and was positive by P/N>2.1.

### 2.12 Westen-Blot detection of anti ACE2 antiserum specificity in rats

After SDS-PAGE electrophoresis of the total tissue protein extracted from the ACE2 fusion protein and the proteins from former, the corresponding strips on the gel were transferred to the PVDF membrane, and the PVDF membrane was placed in the 50 g/mL skimmed milk powder in 2 h at room temperature, and then transferred to the rat anti ACE2 blood albumin diluted with TBST 1:6400 times in overnight at 4°C temperature, and the film was washed with a TBST liquid for 5 times. Then transferred to the Goat anti mouse IgG labeled with 1:5000 diluted horseradish enzyme in TBST solution, incubated at room temperature for 2 h and TBST washing solution for 5 times. After color rendering, the images were observed in the automatic chemiluminescence image analysis system.

### 2.13 Immunofluorescent detection of anti ACE2 serum specificity

The IPEC-J2 cells were cultured in 6 orifice plates with cell crawling slices. After being placed in 37°C and 5%CO_2_ incubators to 70% of fusion degree, After washing with PBS for 2 times, then 4% polyformaldehyde was used to fix 15 min, PBS was washed 5 times and permeated with 0.5%Triton 100 for 20 min, and then washed again, placed in the 50g/mL skimmed milk powder in 2 h at room temperature and then transferred to rat anti ACE2 serum at 4°C incubated overnight. After washing with PBS for 5 times, transferring into the Goat anti rat IgG which diluted by PBS at 1:500 in 2 h at room temperature and washed for 5 times by PBS. The DAPI staining 5 min for 100 mg/mL, and PBS washing for 3 times, observed in a confocal microscope/ultrahigh resolution microscope STORM.

### 2.14 Immunohistochemical identification and localization of murine anti ACE2 protein polyclonal antibody in pig tissues

Immunohistochemical was performed on formalin-fixed tissue. Gut fragments, 3×1 cm (7∼10 cm away from the stomach), were flushed with water, bound in surgical tape and fixed in ice-cold 10% neutral buffered formalin for a maximum of 14 h at 4°C. All fixed samples were embedded in paraffin and sectioned to 5 μm on poly L-lysine slides. Antigen retrieval was performed by boiling in citrate buffer (LabVision, Fremont, California, USA) for 10 min at 100°C. The endogenous peroxidase was blocked (only for immunohistochemical staining) by EnVision blocking solution (DakoCytomation, Glostrup, Denmark) for 5 min. The non-specific binding was blocked for 30 min with 5% rabbit serum (DakoCytomation) for the primary mouse serum(1:600), and primary antibody incubation was performed at 4°C overnight. The immunohistochemical detection was performed using secondary antibody horseradish peroxidase labelled polymer and DAB reagent (Dako Cytomation).

## 3 RESULTS

### 3.1 Cloning and sequence analysis of the target gene of porcine ACE2

4 reverse transcriptional products were randomly selected for PCR amplification. A single band was found in the spleen, duodenum, jejunum and ileum by 1% agarose gel electrophoresis (Line 1-4). The size of the strip was about 2400 bp, and was the same as expected (2418 bp) (Figure 1-A).

**Figure 1.**
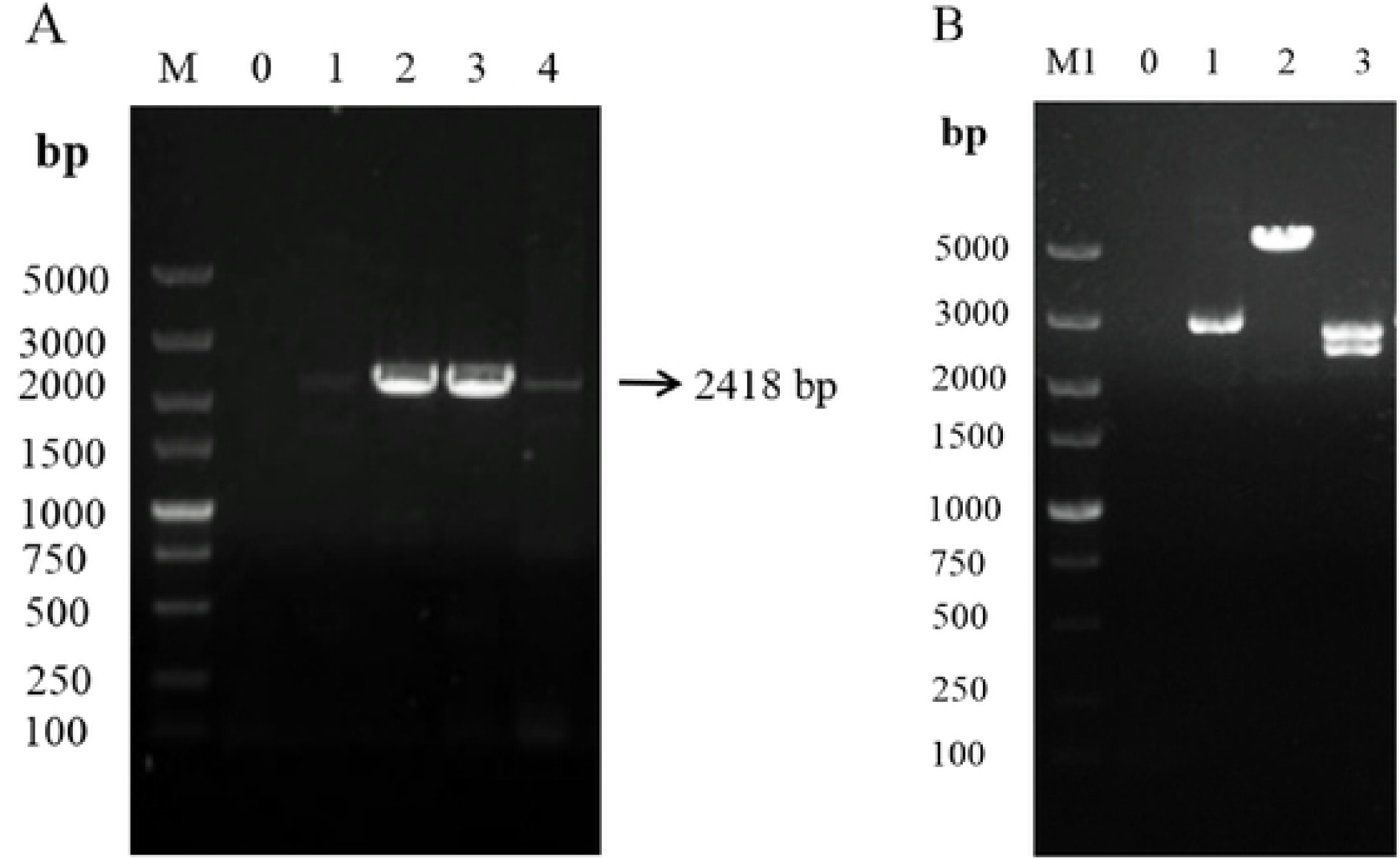
ACE2 gene sequencing. A: The product of PCR amplification of ACE2 gene; M: DL 5000, Line 0: negatively control, Line 1: ACE2 in spleen; Line 2: ACE2 in duodenum; Line 3:ACE2 in jejunum; Line 4: ACE2 in ileum; B: Single and double enzyme digestion of recombinant plasmid. Line 0: negative control; Line 1: Recombinant plasmid ACE2-pMDl9T; Line 2: Recombinant plasmid was digested by Hind 111; Line 3: Recombinant plasmid was digested by BamHI and Sacl.

### 3.2 The recombinant vector of PMD-19T-ACE 2 was successfully constructed

In order to verify whether the ACE2 gene fragment was successfully inserted into the pMD19-T vector, 2 positive colonies were selected and the plasmid was extracted by plasmid DNA, and 1% agarose gel electrophoresis results were found in figure 1-B. As can be seen from figure 1-B, 1 treaty 5110 bp bands were found after the Hind III single enzyme was cut, and the size was consistent with the sum of the pMD-19T vector (2692 bp) and the target gene (2418 bp). BamH I and Sac I double digestion showed about two bands around 2700 bp and 2400 bp, which confirmed that pMD-19T-ACE2 recombinant vector had been successfully constructed. The positive plasmid was selected and sent to Shanghai Ying Jun Biotechnology.

### 3.3 Homology analysis of ACE2

Using DNAstar software, EditSeq template was used to analyze the coding region sequence and homology analysis of wild boar and other genes in Genbank. From Table 1, we can see that the nucleotide sequence obtained from this experiment is compared with the ACE2 nucleotide sequence of 2 pig species found on Genbank, and its homology is 99.5% and 99.1% respectively. Their homology with cattle, goats, domestic cats, domestic dogs, rhesus monkeys, humans and Rattus norvegicus were between 82.0%∼90.0%, and the lowest homology with zebrafish was less than 50%.

**Table 1.** Comparison of nucleic acid homology between the piglet and other organisms

| Species | GenBank accession No. | Homology (%) |
| --- | --- | --- |
| Sus scrofa | NM_001123070.1 | 99.5 |
| Sus scrofa domestica | GQ262781.1 | 99.1 |
| Bos taurus | NM_001024502.4 | 88.8 |
| Felis catus | NM_001039456.1 | 87.4 |
| Canis lupus familiaris | NM_001165260.1 | 86.6 |
| Rattus norvegicus | NM_001012006.1 | 82.0 |
| Macaca mulatta | FJ170098.1 | 84.8 |
| Capra hircus | NM_001290107.1 | 89.2 |
| Danio rerio | NM_001007297.1 | 49.7 |
| Homo sapiens | NM_021804.2 | 84.9 |

According to the results of nucleotide sequencing, the corresponding amino acid sequence was conjectured. The piglet encodes 805 amino acids, which is consistent with the number of ACE2 amino acids in human and Rattus norvegicus, and an amino acid difference with cattle and sheep (804 amino acids).

### 3.4 Homology analysis of ACE2

Using the MegAlign module in DNAstar software, the amino acid sequences derived from the cloned pig ACE2 nucleotide sequence and the amino acid sequences derived from wild boar and domestic pig ACE2 nucleotides were 99.3% and 98.5%, respectively, compared with the other species ACE2 amino acid sequences, and the results were compared with sheep, mountain cattle, domestic cats, domestic dogs, macaques, brown, brown, and brown. Homology of rodents, humans and alpacas reached over 80%, and the lowest homology with zebrafish was 46.8% (Fig. 2-A). From the tree of genetic evolution (Figure 52-B), the fragment is in the same evolutionary group with the ACE2 amino acid sequence of goats and cattle, and the zebrafish is in a completely different 2 branch, and the genetic evolution is consistent with the evolution of amino acid.

**Figure 2.**
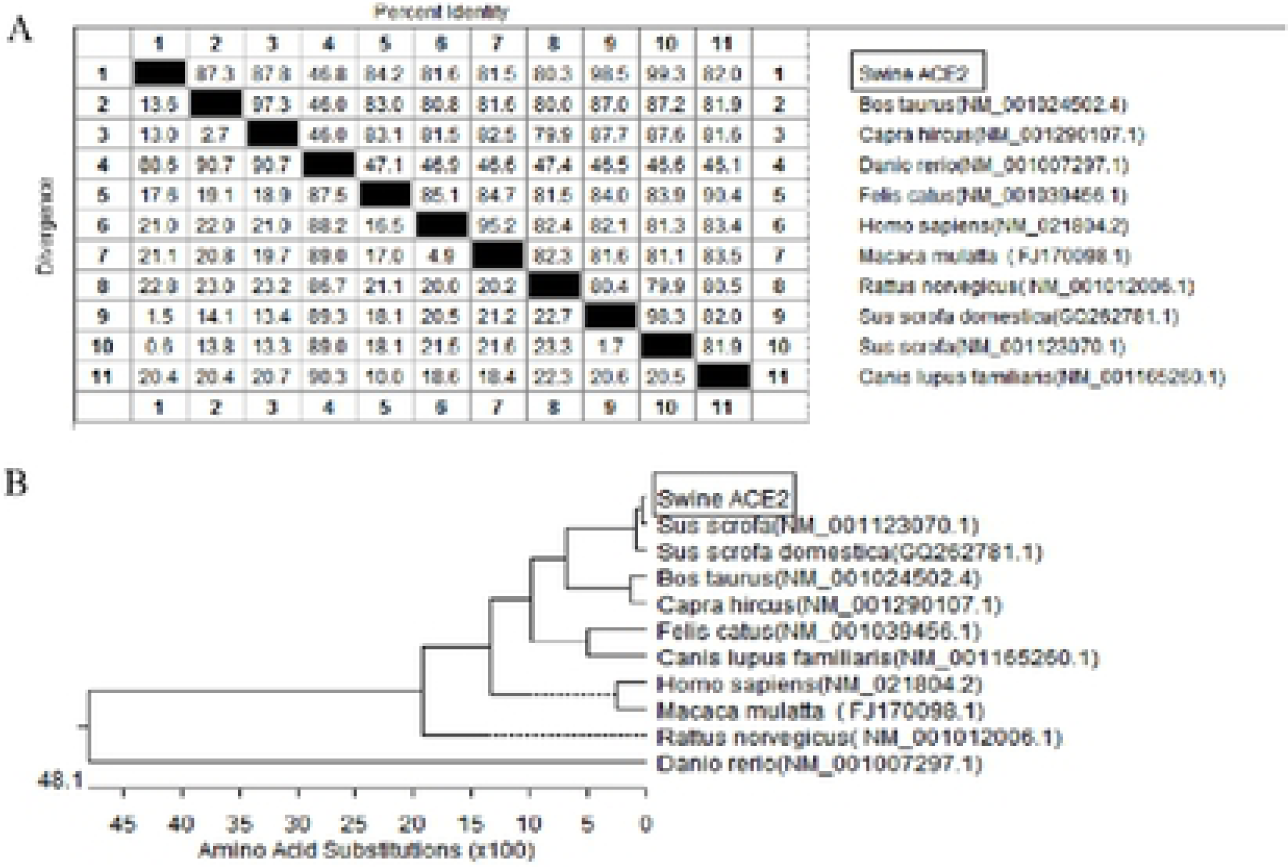
Homology / phylogenetic tree analysis of ACE2 protein. A: Comparison of the deduced amino acid sequences betweendifferent species of ACE2; B: Phylogenetic tree of the deduced amino acid sequence of ACE2 nucleotide sequences between d ifferent species.

### 3.5 Structural Analysis of ACE2 protein

The cross membrane region analysis of weaned piglets ACE2 protein was analyzed by TMHMM2.0 software. The results showed that the porcine ACE2 protein belonged to I type transmembrane protein, and the transmembrane types of ACE2 proteins, such as people and sheep, were the same. Among them, part 1∼739aa is the extracellular part, and there is a transmembrane helix between 740∼762 aa, and the 763∼805 AA is the intracellular part (Fig. 3-A).

**Figure 3.**
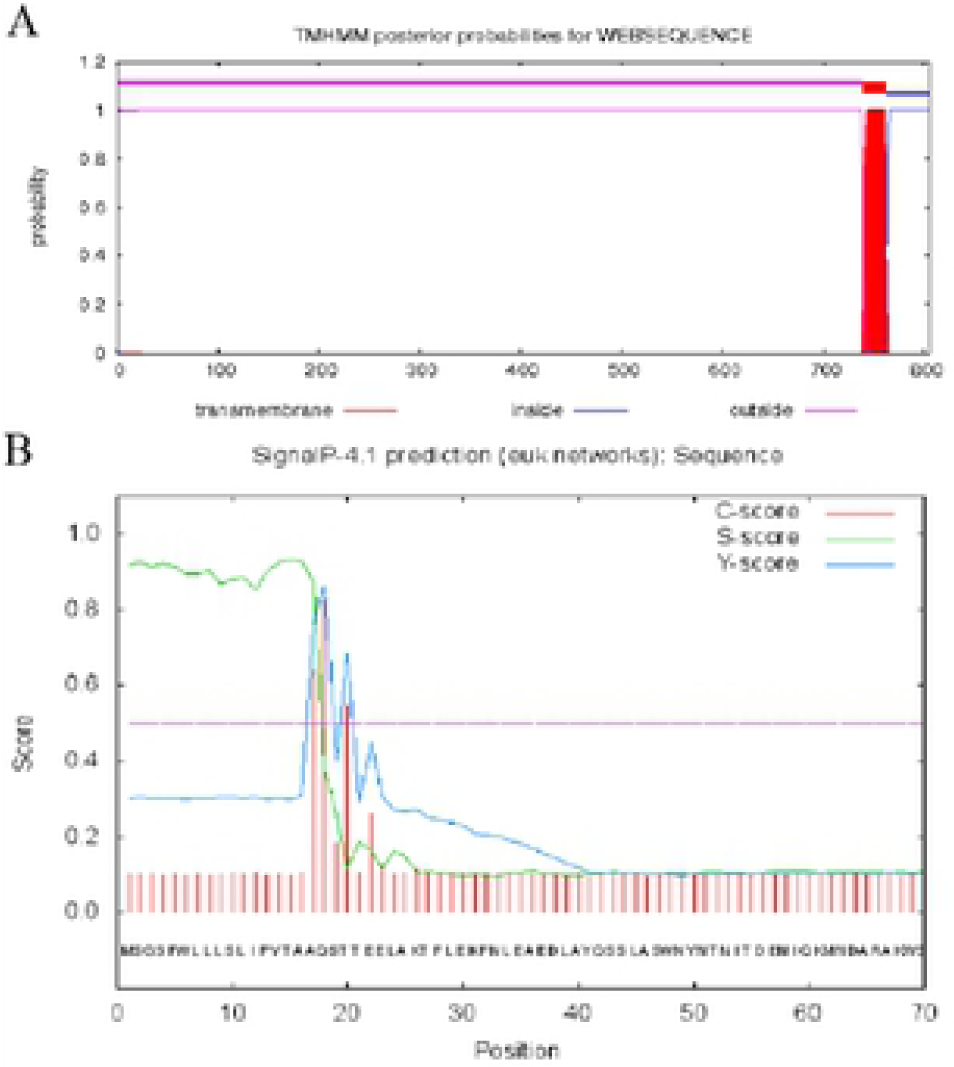
Prediction of transmembrane domain /signal peptide analysis of ACE2 protein from piglet. A: Prediction of transmembrane domain; B: signal peptide analysis.

The Signal P4.1 software predicts the amino acids encoded by the pig ACE2 gene. The results are as shown in figure 3-B: the C value at the frame is maximum, the S value is steep, and the Y value is the highest. It is predicted here that there is a potential cracking site between 17-18aa and the protein is secreted protein (this is judged by the value of D, D is the average value of S-mean and Y-max. It is important to distinguish whether the protein is secreted or not, the D value is 0.883, greater than that of 0.450--, which is the criterion for the secretion of protein. Through online SMART analysis (http://smart.embl-heidelberg.de/), the signal peptide region is between 1-17aa, which is consistent with the results predicted by the transmembrane region analysis.

### 3.6 ACE2 protein has good hydrophilicity

The hydrophilicity of ACE2 protein was analyzed according to the amino acid hydrophilic standard of Kyke-Doolittle. Results are shown in Figure 4. The proportion of hydrophilic region of ACE2 protein in piglets is larger, and the distribution is more uniform (more than 0 lines in the map), indicating that the protein has good hydrophilicity.

**Figure 4.**
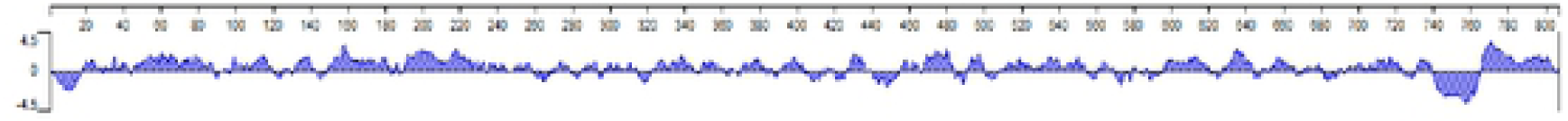
Hydrophilic analysis of ACE2 protein in piglet.

**Figure 5.**
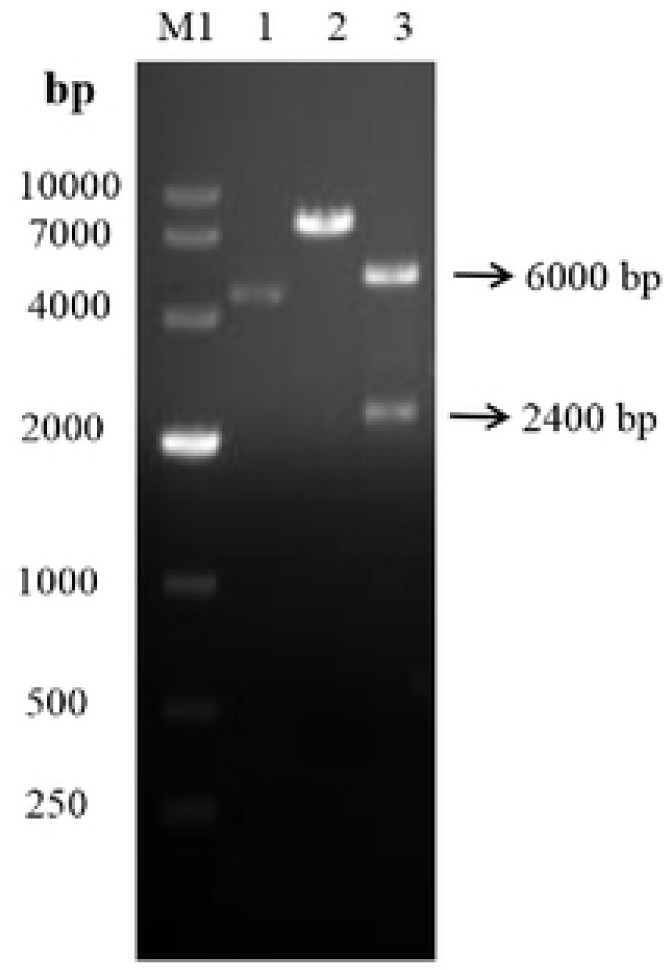
Development of prokaryotic expression construction of ACE2. M_1:_ DLI0000; Line 1: Recombinant plasmid; Line 2: Recombinant plasmid was digested by Hind III; Line 3: Recombinant plasmid was digested by BarnHI and SacI.

### 3.7 Construction and identification of prokaryotic expression vectors

After the recombinant plasmid and the expression vector were cut by BamH I and Sac I, the T4 DNA ligase was linked and transformed into E.coli BL21 (DE3), and the positive colony was picked up for PCR, and then the plasmid was identified by BamH I and double enzyme digestion. 1% agarose gel was used to detect the strain and about two bands. (Fig.5) the sequencing results were correct, indicating that pET-32a-ACE2 recombinant plasmid was successfully constructed.

### 3.8 Induced expression, identification and purification of recombinant protein

The recombinant expression bacteria were induced and expressed at the final concentration of IPTG at 1 mmol/L. The expression products were collected at different time points. After SDS-PAGE analysis, the target protein was obtained in the induction of 2, 4, 6, 8, 10 h and overnight, and the expression of target protein was the highest when 10 h was induced (Figure 6-A). After ultrasonic breakage centrifugation, the supernatant and precipitation were taken respectively. After SDS-PAGE, the recombinant protein of ACE2 was found mainly in the precipitate, that is, in the form of inclusion body (Figure 6-B). The purified fusion protein (Fig 6-C) was purified and purified by KCl staining, which could be used for subsequent polyclonal antibody preparation.

**Figure 6.**
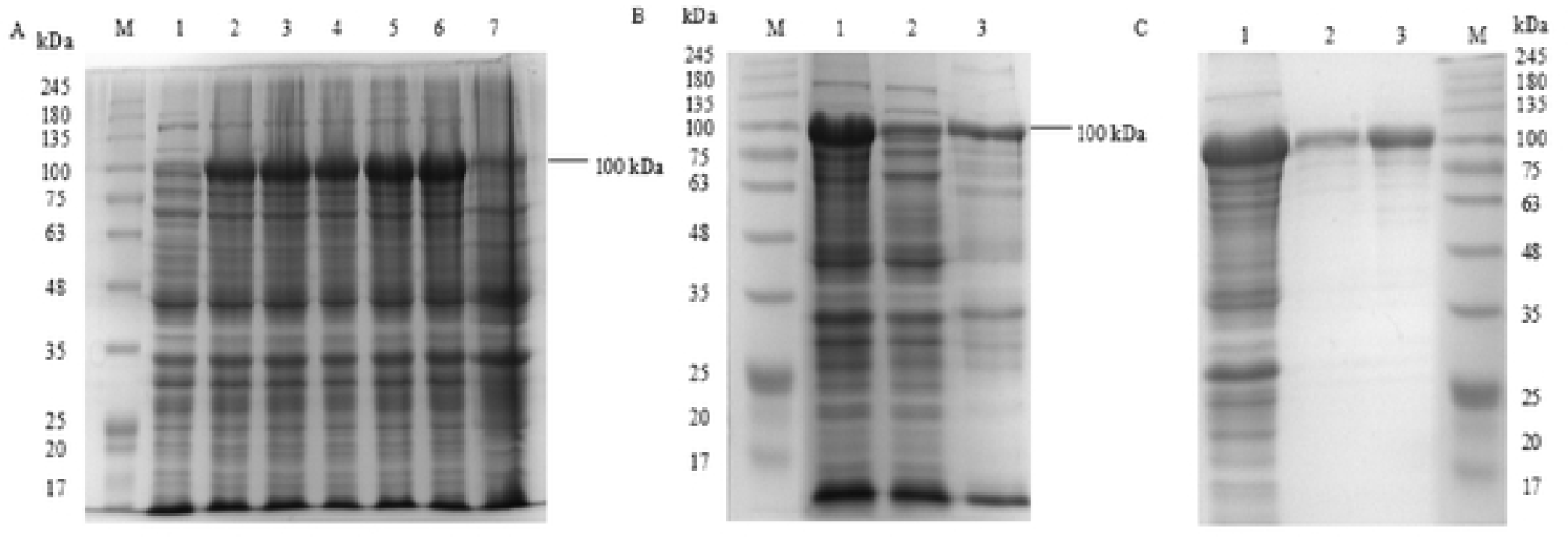
Expression Purification of pET-32a-ACE2 fusion protein. A: Induced expression of pET-32a-ACE2 in E:coli BL21(DE3); M: Protein molecular weight Marker; Line 1∼7: Recombinant pET-32a-ACE2 expression products in E:coli BL2l(DE3) induced with IPTG at 0, 2, 4, 6, 8, 10 and overnight; B: Solubility analysis ofpET-32a-ACE2 fusion protein; M: Protein molecular weight Marker; Line 1: Recombinant bacterium; Line 2: The upernatant; Line 3: The deposit; C: Purification of pET-32a-ACE2 fusion protein; M: Protein molecular weight Marker; Line 1 : Recombinant bacterium; Line 2∼3: Purificated fusion protein.

### 3.9 Determination of anti ACE2 polyclonal antibody titer by indirect ELISA method

The purified pET-32a-ACE2 fusion protein was immunized with 4 Wastar female rats. After 4 immunizations, the antibody test results were all positive, the rats were slaughtered, the isolated serum obtained the anti ACE2 antibody of the polyclonal mice, and the antiserum titer was detected by indirect ELISA method. Results as shown in Figure 7, the serum dilution ratio of the positive serum OD_450_ was about 1, and the P/N value (P: the mean OD_450_ value of the positive hole, the mean value of the negative pore, OD_450_) was the best serum dilution degree, that was 3200.

**Figure 7.**
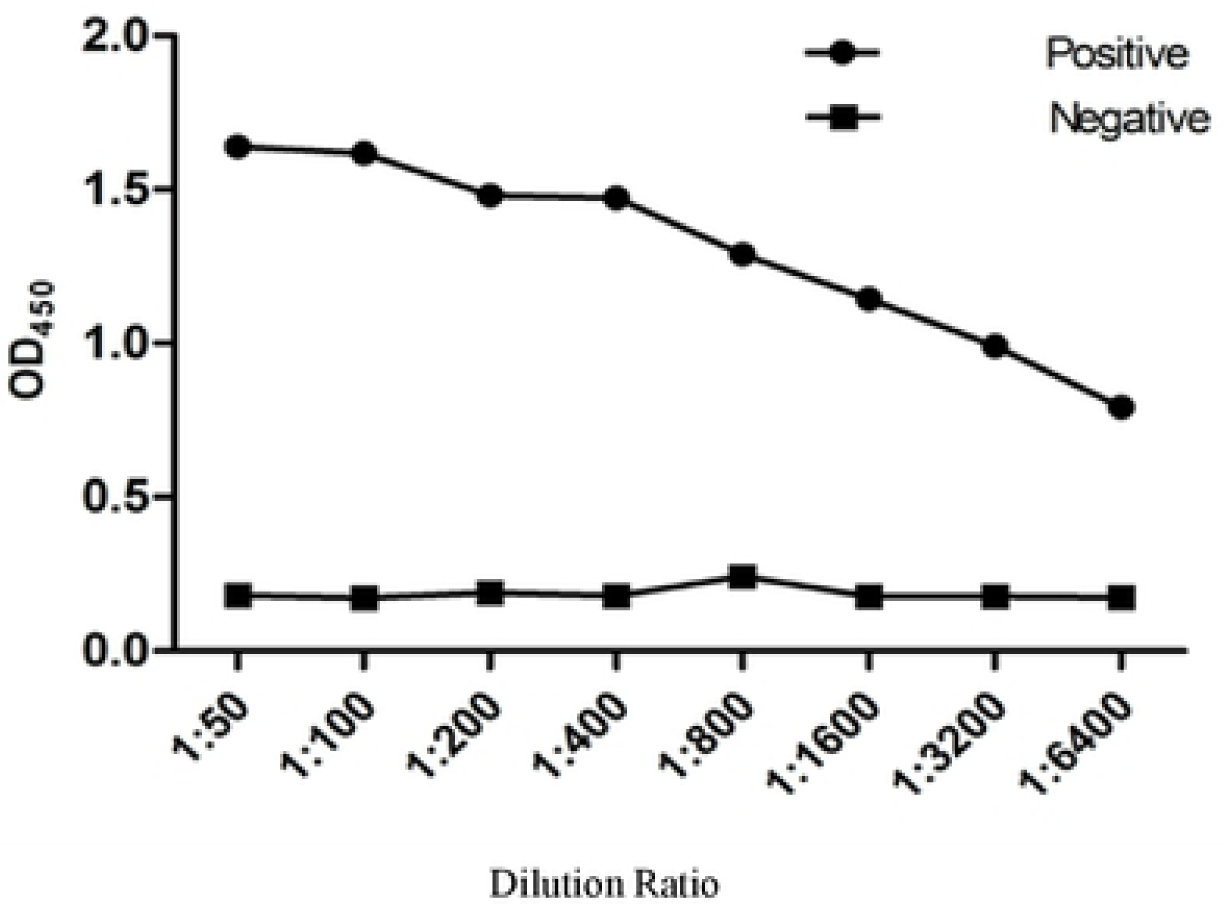
ELISA detection of rats anti-ACE2 titer.

### 3.10 Detection of serum specificity of anti ACE2 in rats by Western blot

The rat anti ACE2 serum was prepared with the best multiple dilution as a single antibody, and the Sheep anti rat IgG-HRP was used as a two antibody. The purified fusion protein, pig heart, spleen, kidney, liver, duodenum, sheep liver, rat liver, rat liver and chicken liver and kidney tissue protein were analyzed by Western blot. The results showed the fusion protein and pig heart. There was a specific binding protein in the size of 100 kDa in the viscera, spleen, kidney, liver and duodenum (Fig. 8), and the rest did not appear. The results show that the ACE2 polyclonal antibody can identify the fusion protein and the ACE2 protein in the pig tissues, but the ACE2 protein in other species can not be combined, that is, the ACE2 polyclonal antibody against the pig’s mouse has good immune response and specificity.

**Figure 8.**
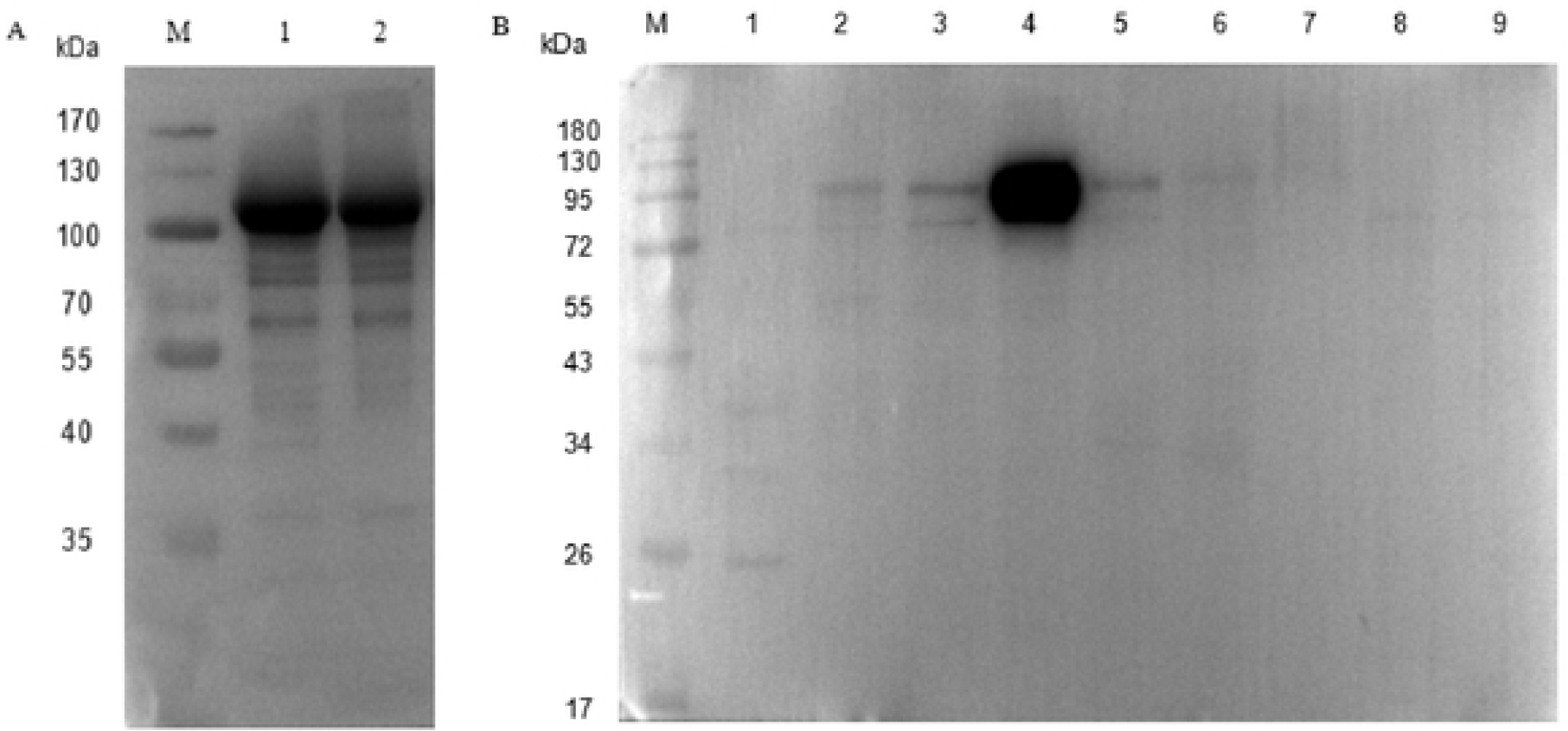
Westernblotting analysis of specificity of polyclonal antibody. A: Western blotting analysis of purified protein; M: Protein molecular weight Marker; Line 1∼2: purified protein; B: Western blotting analysis of purified protein of different tissue proteins; M: Protein molecular weight Marker; Line 1: Cardiac tissue protein of porcine; Line 2: Spleen tissue protein of porcine; Line 3: Duodenum tissue protein of porcine; Line 4: Kidney tissue protein ofporeine; Line 5: Liver tissue protein of porcine; Line 6: Liver ti ssue protein of goat; Line 7: Liver tissue protein of rat; Line 8: Liver ti ssue protein of chicken; Line 9: Kidney tissue protein of chicken.

### 3.11 Anti ACE2 serum specificity in mice by immunofluorescent

The mouse anti ACE2 serum was diluted by multiple 1:600 as one antibody, the fluorescent enzyme labeled Sheep anti mouse IgG-HRP was two resistance, and the final concentration was 100 ng/mL DAPI for 5 minutes. The immunofluorescence analysis of the pig intestinal epithelial cells, as shown in Figure 9, that the mouse anti ACE2 could specifically identify the ACE on the membrane of the pig small intestinal epithelial cells. The 2 protein, that is, the polyclonal antibody of pig anti mouse ACE2 prepared, has better immunoreactivity.

**Figure 9.**
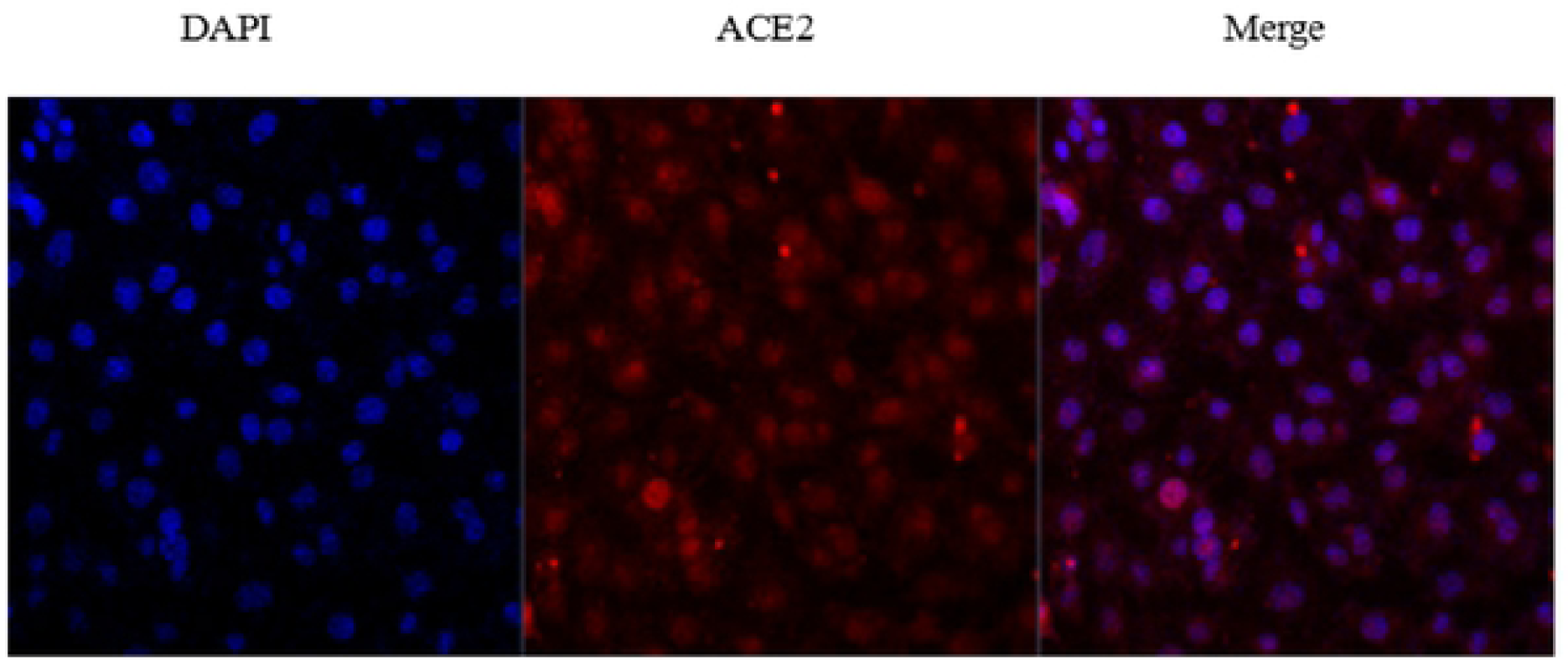
Immunofluorescence analysis of specificity of polyclonal antibody.

### 3.12 Serum sensitivity of anti ACE2 in rats by Western blot

As shown in Figure 10, the proteins extracted from pig spleen, kidney, liver, lung, pancreas, lymph node, stomach, duodenum, jejunum, ileum, colon, rectum and cecum were analyzed by Western blotting, and the rat anti porcine polyclonal antibody was diluted with 1: 3200 as one resistance, and the IgG of sheep resistant to HRP was two. Resistance, the results showed that the tissue protein extracted from pig kidney, jejunum and rectum had obvious bands at 92 kDa, and the highest content in kidney, but not in other tissues. Under the same condition, the Sheep anti rabbit polyclonal antibody was diluted with 1:1000 multiple as a single antibody, and the HRP labeled Goat anti rabbit IgG was two. The results showed that the tissue protein extracted from pig spleen, kidney, lung, pancreas, lymph node, stomach, duodenum, jejunum, ileum, colon, rectum, and cecum was marked at 92 kDa. The results showed that the sensitivity of mouse against polyclonal antibody was poor.

**Figure 10.**
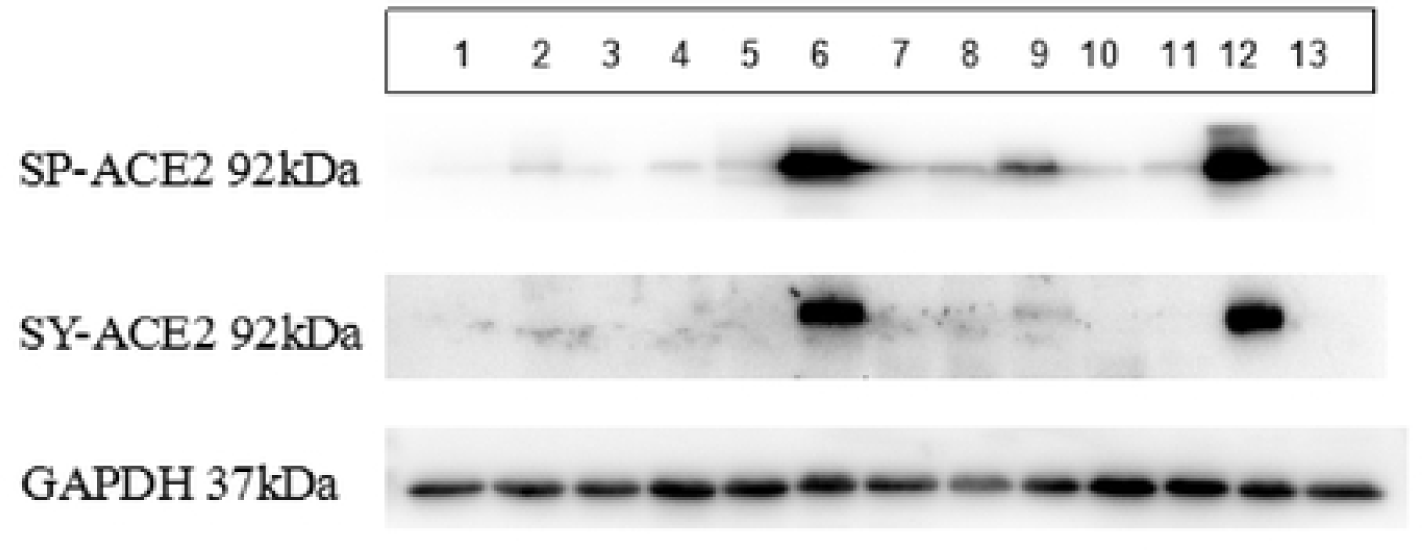
Western blotting analysis of differert tissues from porcine. Line 1: Live tissue protein of porcine; Line 2: Spleen tissue protein of porcine; Line 3: lymph tissue protein of porcine; Linc 4: stomach tissue protein of porcine; Line 5: duodenum tissue protein of porcine; Line 6: jejunum tissue protein of porcine; Line 7: ileum tissue protein of porcine; Line 8: colon tissue protein of porcine; Line 9: rectum tissue protein of porcine; Line 10: caecum tissue protein of porcine; Linc 11: lung tissue protein of porcine; Line 12: kindcy tissue protein of porcine; Line 13: pancreas tissue protein of porcine.

### 3.13 Immunohistochemistry was used to detect the expression of ACE2 in pig tissues

The cellular localization of ACE2 in porcine tissues was analyzed after dilution of 1:600 prepared by anti ACE2 serum. The criterion is that if Brown is positive, the result is shown in Fig. 11. As can be seen from figure 3-21, ACE2 protein exists in various tissues and organs of the pig, mainly expressed in the mucosa of the gastric fundus (Fig. 11-A), the myometrium of the small intestine, serous layer, and villi epithelium (FIGS. 11-B, C, D). In the large intestine, it is mainly expressed in the intestinal gland (FIGS. 11-E, F), and the result of this group on the pig WB Like. The results further confirmed the specificity of the mouse anti ACE2 polyclonal antibody prepared in this experiment.

**Figure 11.**
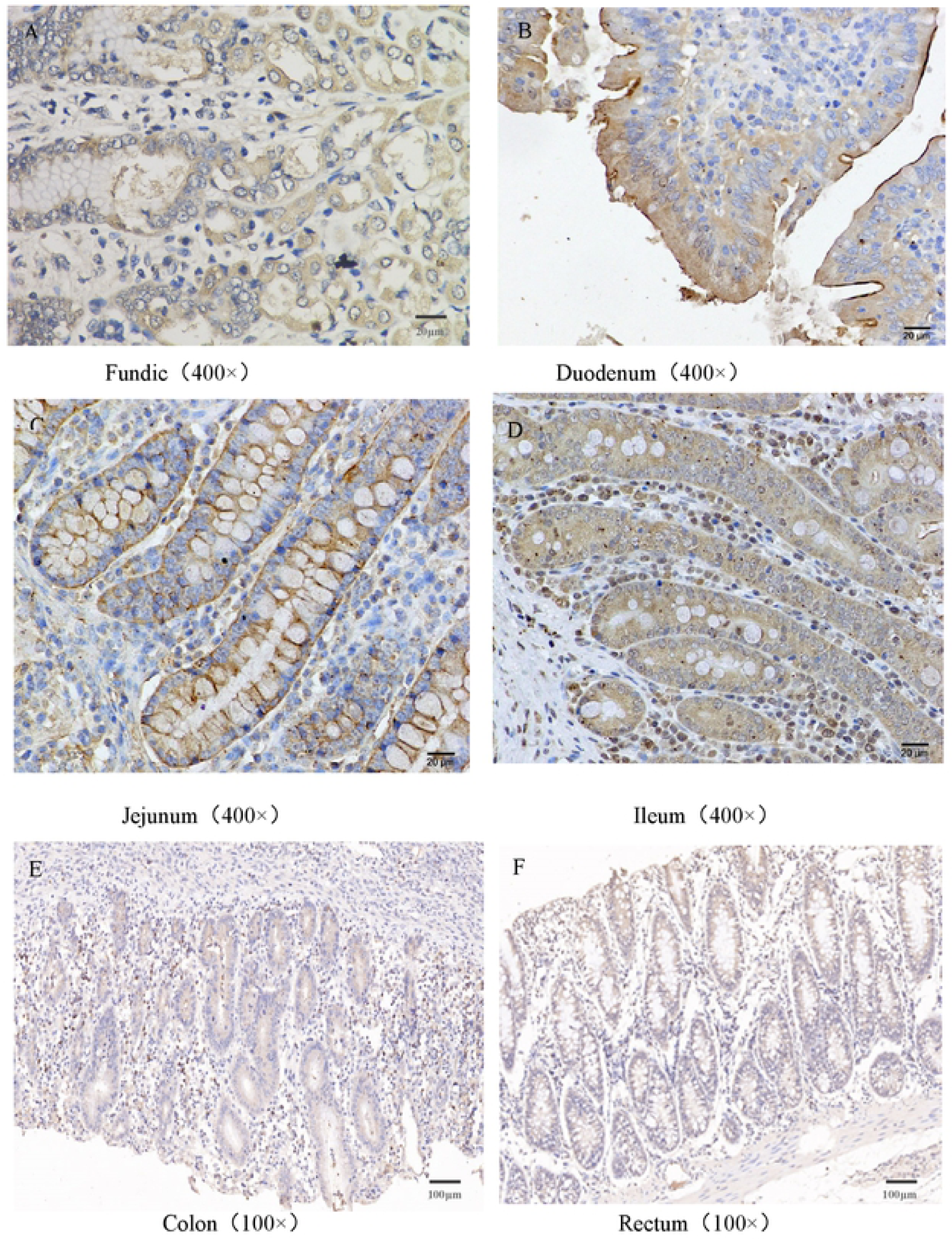
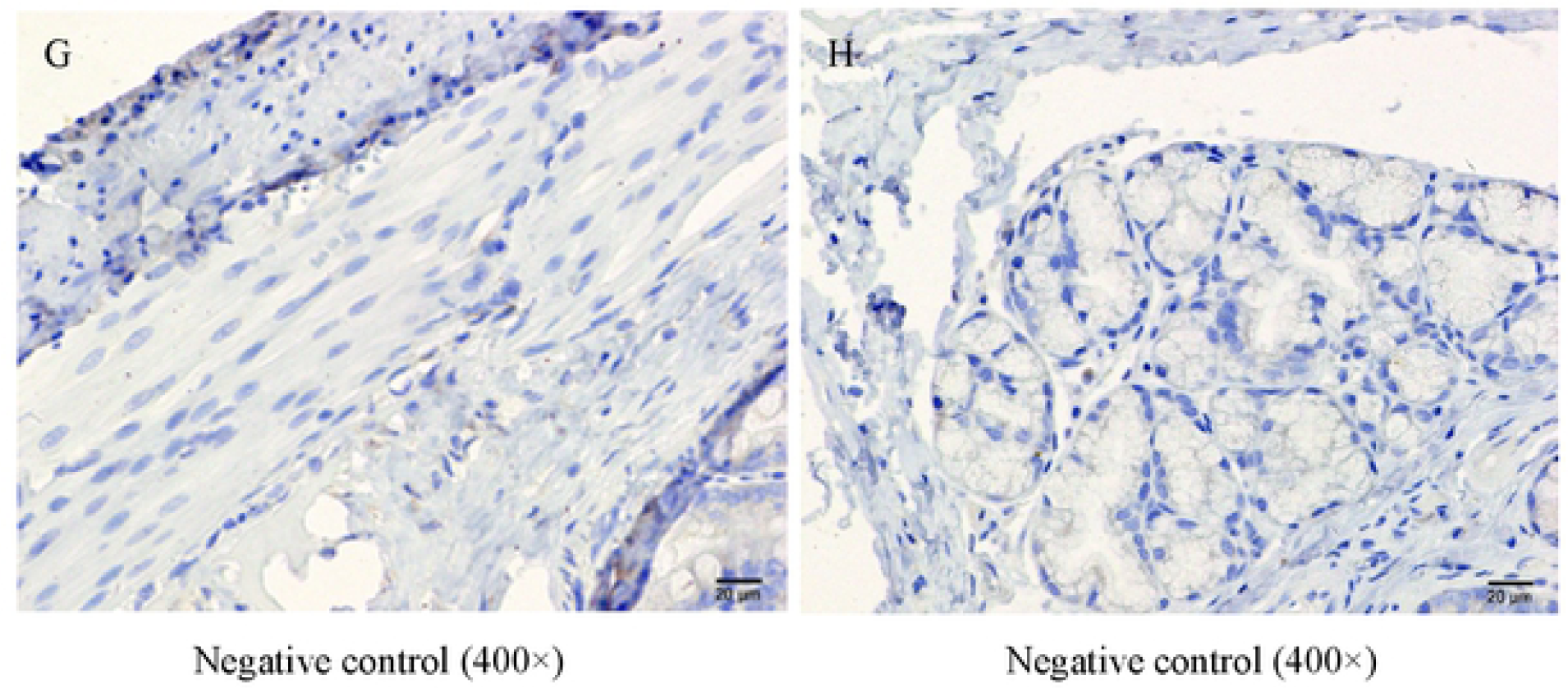
ACE2 immunohistochemical map of differert tissues from pigs.

## 4 DISCUSSION

In 2000, Tipnis amplified the genes encoding ACE2 from the cDNA Library of human lymphoid cancer(Tipnis et al., 2000). It was named as ACEH and numbered as AF241254 in the gene pool. Further study found that the open reading frame of cDNA of the ACEH was composed of 2418 nucleotides, encoding 805 amino acids, and the gene was located at the chromosome Xp22 site and contains 18 exons. In the same year, Donoghue et.al constructed a full-length ACE2 expression vector from the cDNA Library of heart failure patients through transfection of CHO cells(Donoghue et al,. 2000). The gene sequence was recorded by GenBank and numbered as AF291820. Later, it was confirmed that ACE2 and ACEH were the same. In 2003, Moore confirmed that ACE2 was the main functional receptor of the SARS coronavirus(Li et al,. 2003). In order to further understand the route of infection, Li cloned the full-length ACE2 sequence of the beaver, and confirmed that the beaver was infected mainly by the S protein region of the SARS coronavirus combined with its ACE2(Ge et al,. 2013; Li et al,. 2005). Wang extracted the total RNA from the lungs of 9 month old cats and amplified the ACE2 gene sequence of the home cat by RT-PCR(Wang et al. 2005). The gene number in Genebank was AY957464 and its sequence size was 2418 bp. It encodes 805 amino acids. It was suggested that cat ACE2 might be more likely to mediate the invasion of SARS-CoV. In 2005, Xie successfully amplified the full length cDNA sequence of the ACE2 gene from the mouse kidney tissue, and the sequence size was 2418 bp(Xie et al,. 2005). Then Xu uploaded the partial ACE2 gene sequence of kidney of Trionyx sinensis(Genbank: HM107424)(Xu et al,. 2011). In 2018 Chen cloned the whole ACE2 gene sequence from the kidney and lung of monkeys. Compared with the human ACE2 sequence, there were 38 NS mutations(Chen et al,. 2008). The specific fragment of the ACE2 gene coding region of the goat kidney tissue was amplified by Yang in the laboratory for the first time. The length of the nucleotide sequence was 2415 bp(Yang et al,. 2016).

In this study, the pig ACE2 gene was cloned, analyzed and prokaryotic expressed successfully, and the polyclonal antibody of mouse ACE2 was prepared. The whole gene sequence of ACE2 in pig was 2418 bp and 805 amino acids were encoded. Homology analysis on GenBank found that the cloned ACE2 gene were above 99% similar to other porcine ACE2 genes, which shows that the pig ACE2 gene cloning was successful. The full-length gene sequence of pig ACE2 gene was provided to GenBank database., It is deduced that the pig ACE2 protein belongs to the type I secretory transmembrane protein and its signal peptide sequence is located between 1∼17 AA, which is similar to that of goats (1∼18 AA) (Yang et al,. 2016).

In conclusion, In this paper, the basic information of porcine ACE2 protein was provided and the specific anti-porcine polyclonal antibody was successfully, which could lay the foundation for further research on the biological activity of ACE2 in pigs.

## ACKNOWLEDGMENTS

This work was supported by the National Natural Science Foundation of China (31972640).

